# A subtle structural modification of a synthetic cannabinoid receptor agonist drastically increases its efficacy at the CB1 receptor

**DOI:** 10.1101/2023.06.10.544442

**Authors:** Hideaki Yano, Rezvan Chitsazi, Christopher Lucaj, Phuong Tran, Alexander F. Hoffman, Michael H. Baumann, Carl R. Lupica, Lei Shi

## Abstract

The emergence of synthetic cannabinoid receptor agonists (SCRAs) as illicit psychoactive substances has posed considerable public health risks that include fatalities. Many SCRAs exhibit much higher efficacy and potency, compared with the phytocannabinoid Δ^9^-tetrahydrocannabinol (THC), at the cannabinoid receptor 1 (CB1R), a G protein-coupled receptor involved in modulating neurotransmitter release. In this study, we investigated structure activity relationships (SAR) of aminoalkylindole SCRAs at CB1Rs, focusing on 5F-pentylindoles containing an amide linker attached to different head moieties. Using *in vitro* bioluminescence resonance energy transfer (BRET) assays, we identified a few of SCRAs exhibiting significantly higher efficacy in engaging the Gi protein and recruiting β-arrestin than the reference CB1R full agonist CP55940. Importantly, adding a methyl group at the head moiety of 5F-MMB-PICA yielded 5F-MDMB-PICA, an agonist exhibiting a large increase in efficacy and potency at the CB1R. This pharmacological observation was supported by a functional assay of the effects of these SCRAs on glutamate field potentials recorded in hippocampal slices. Molecular modeling and simulations of the CB1R bound with either of the SCRAs revealed critical structural determinants contributing to the higher efficacy of 5F-MDMB-PICA, and how these subtle differences propagated to the receptor-G protein interface. Thus, we find that apparently minor structural changes in the head moiety of SCRAs can cause major changes in efficacy. Our results highlight the need for close monitoring of structural modifications of newly emerging SCRAs and their potential for toxic drug responses in humans.

## INTRODUCTION

Synthetic cannabinoid receptor agonists (SCRAs) were originally developed as research compounds for studying the endocannabinoid system and subsequently repurposed as nausea suppressants, appetite stimulants, and for neuropathic pain relief [1, 2]. Unfortunately, since their introduction in medicinal chemistry, SCRAs have extensively infiltrated the recreational drug market as new psychoactive substances (NPS), leading to significant public health threat due to their elevated potential to cause psychiatric and medically serious disorders as well as mortality in humans. Over the past decade, the relatively widespread use of these unregulated NPS has led to numerous cases of mass overdoses, accidental intoxications, and fatalities [3–5]. Moreover, the increased efficacy and potency of SCRAs synthesized in illicit laboratories is associated to increased risks of their adverse and toxic effects [6]. A striking example of the danger posed by SCRAs is a mass intoxication event that occurred in New York City in 2016, commonly referred to as a “zombie outbreak”. This event resulted from the use of the molecule MMB-FUBINACA, which resulted in symptoms including catatonia, seizures, and psychosis [4]. Since this event, numerous additional reports have emerged describing deaths from the use of 5F-MDMB-PINACA [3, 7]. Where investigated, it was observed that these SCRAs exhibit high potency at CB1Rs and CB2Rs. Additionally, these drugs act as full agonists at CB1Rs, which distinguishes them from endogenous cannabinoids and the well-known psychoactive constituent of cannabis, THC [8–10].

The CB1R is one of the most abundant G protein-coupled receptors in the brain that primarily activates Gi/o proteins to inhibit adenylyl cyclases to inhibit cAMP production leading to the inhibition of gene transcription and synaptic remodeling [11–13]. CB1Rs are widely expressed on axon terminals where they can be activated by endocannabinoids. CB1Rs can also activate G protein-coupled inwardly-rectifying potassium channels (GIRKs) to hyperpolarize neurons and inhibit voltage-gated calcium channels (VGCCs), primarily through liberation of G protein β and γ subunits [14–17]. The inhibition of neurotransmitter release by CB1Rs has been identified in many neuronal pathways, including neurotransmission mediated by glutamate and GABA, acetylcholine, and monoamines, such as serotonin [14–17]. In addition, CB1Rs expressed on astrocytes are reported to regulate the release of gliotransmitters that can further modulate neuronal activity [18, 19]. Together, through their ability to suppress neurotransmission in many neuronal circuits, CB1Rs and the endocannabinoid system regulate a wide variety of physiological functions, including learning and memory, motivation, pain, and anxiety [10, 12].

It is noteworthy that SCRAs induce a distinct subset of adverse effects such as nausea, anxiety, hypothermia, hallucination, catalepsy, tachycardia, hypertension, and withdrawal, which are unlike those associated with phytocannabinoids [20–24]. Many SCRAs share an aminoalkylindole scaffold, which is divided into four connected moieties known as the “head, linker, core, and tail” [1]. Structure-activity studies have identified several high efficacy agonists at the CB1R, including 4CN-CUMYL-BUTINACA, 5F-MMB-PINACA, and 5F-CUMYL-PINACA, among others. All of these exhibit much higher potency and efficacy at CB1Rs, compared to phytocannabinoids and many first generation SCRAs, and this has been associated with much higher incidences of neurological symptomology [25]. Additionally, some adverse effects of these SCRAs may also be attributed to off-site interactions with non-canonical receptor targets [26].

Crystallographic and cryo-EM studies of CB1Rs have greatly advanced our mechanistic understanding of their function, revealing both active and inactive conformational states at the atomistic level [27–32]. Notably, the active molecular structures bind to a variety of CB1R agonists, not only the widely used reference SCRA CP55940, but also a few other SCRAs such as AM11542, AM841, and MDMB-FUBINACA. Importantly, the active CB1R structures in complex with Gi protein can serve as excellent templates for investigating the SAR of SCRAs.

Synthetic orthosteric agonists that share binding sites yet elicit a higher maximum response than the endogenous agonist, resulting in E_max_ greater 100%, are considered as superagonists for the target receptor [33, 34]. However, this concept is often contentious as the readouts used to compare efficacies may be influenced by signal amplification, which leads to higher apparent E_max_ values. Despite the complexities in assay interpretation and subject receptor system, there are instances where synthetic ligands confer superagonism [35–37].

5F-pentylindoles, a subgroup of highly potent and efficacious SCRAs, have been associated with a surge in human use and coinciding hospital visits [24]. In this study, we aim to investigate 5F-pentylindoles and analyze their analogues especially within their “head” moiety that manifests as efficacy above the full agonist. Here, we provide molecular mechanisms that may explain superagonism at the CB1R.

## RESULTS

### Subtle changes in the head moiety of 5F-pentylindoles result in distinct pharmacological profiles of Gi1 engagement

Aminoalkylindole SCRAs, such as JWH018 and AM2201, which exhibit high potency and efficacy at CB1Rs, were widely abused in early 2010’s [38–40]. In this study, we investigated the SAR of aminoalkylindole SCRAs specifically with a 5F-pentylindole scaffold and focused on several molecules containing an amide linker attached to different head moieties. To study CB1R-G_i1_ coupling, we used G protein engagement BRET in heterologous HEK293 cells [41]. This method has the advantage over other functional assays such as adenylyl cyclase inhibition, as it provides a direct measurement of G_i1_ coupling and is not confounded by potential off-target receptor activation of SCRAs. We collected readouts at four time points (2, 16, 30, 44 min) of ligand incubation to capture kinetic profiles of receptor G-protein coupling, as lipophilic ligands are known to have binding kinetics different from hydrophilic ones [42, 43]. Results obtained at the 16-min time point were utilized in the analysis as the responses of most agonists reached a stable plateau by this time point that did not change.

Our results show that AM2201 displays a similar potency and efficacy profile to a commonly used reference full agonist CP55940, with a slightly higher E_max_, although the difference is not significant (Table 1). Conversion of the ketone moiety of AM2201 to an amide linker results in a less potent compound (*N*-1-naphthyl, 5F-NNEI) compared to AM2201 (1-naphthyl) (see Figure 1, Supplementary Figure 1, Table 1). Substitution of 1-naphthyl of 5F-NNEI with a less bulky benzyl group (*N*-1-benzyl, 5F-SDB-006) substantially lowers the efficacy but slightly increases the potency relative to 5F-NNEI.

**Figure 1.**
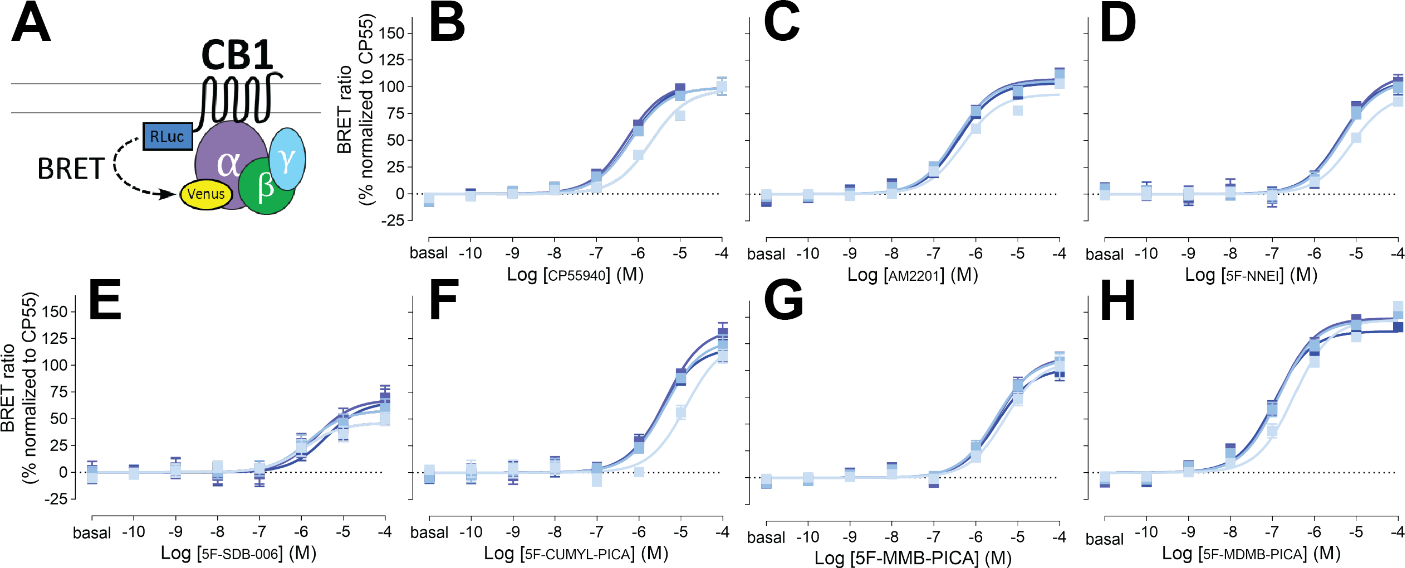
G_i1_ engagement BRET. Drug-induced engagement BRET between CB1R-Rluc and Gi1-Venus (A) measured in response to CP55940 (B), AM2201 (C), 5F-NNEI (D), 5F-SDB-006 (E), 5F-CUMYL-PICA (F), 5F-MMB-PICA (G), 5F-MDMB-PICA (H) at various time points (2, 16, 30, 44 min light to dark blue). Concentration-response curves are plotted as a percentage of maximal response by CP55940 at each time point and presented as means ± SEM of *n* ≥ 3 independent experiments.

**Table 1.** Pharmacological comparison of 5-fluoropentylindole ligands at CB1R. Mean ± SEM values for E_max_ and pEC_50_ are reported. E_max_ is normalized against CP55940. ^#^ Y-axis cut-off value at the highest concentration is shown. ND – EC_50_ cannot be determined. One-way ANOVA followed by Tukey test *p<0.05, ****p<0.0001 against corresponding CP55940 in the same assay.

CB1R-Gi1 engagement at 16 min
| | CP55940 | AM2201 | 5F-NNEI | 5F-SDB-006 | 5F-CUMYL-PICA | 5F-MMB-PICA | 5F-MDMB-PICA | WIN55-212-2 | $\Delta^9$-THC | AEA | 2AG |
| --- | --- | --- | --- | --- | --- | --- | --- | --- | --- | --- | --- |
| E<sub>max</sub> (%) | 100.0±3.2 | 105.8±2.3 | 106.2±6.5 | 57.9±6.2**** | 124.2±6.7* | 111.2±5.2 | 142.1±2.7**** | 98.8±5.2 | 36.1±4.1****# | 57.8±3.1****# | 61.2±2.7****# |
| pEC<sub>50</sub> (M) | 6.19±0.08 | 6.47±0.06 | 5.33±0.12* | 5.83±0.25 | 5.38±0.11 | 5.53±0.10 | 6.85±0.06 | 5.00±0.09**** | ND | 4.92±0.49 | 5.21±0.16 |

CB1R- $\beta$ arrestin2 recruitment at 16 min
| | CP55940 | AM2201 | 5F-NNEI | 5F-SDB-006 | 5F-CUMYL-PICA | 5F-MMB-PICA | 5F-MDMB-PICA | WIN55-212-2 | $\Delta^9$-THC | AEA | 2AG |
| --- | --- | --- | --- | --- | --- | --- | --- | --- | --- | --- | --- |
| E<sub>max</sub> (%) | 100.0±3.1 | 106.5±1.9 | 110.3±2.9 | 46.0±2.5**** | 91.0±3.8 | 99.9±4.1 | 147.7±1.6**** | 121.3±4.7# | 36.0±7.1****# | 49.9±3.0****# | 85.3±4.9# |
| pEC<sub>50</sub> (M) | 5.19±0.06 | 6.12±0.04* | 5.28±0.05 | 5.61±0.12 | 5.86±0.10* | 5.44±0.09 | 6.24±0.03* | 4.72±0.07 | ND | 4.95±0.11 | 4.55±0.92 |

Replacing the benzyl group of 5F-SDB-006 with a cumyl group, which effectively adds two methyl groups and results in 5F-CUMYL-PICA, substantially increased E_max_ to 124% of that of the reference CP55940. Substituting the benzyl group of 5F-SDB-006 with a methyl 3-methylbutanoate group, resulting in 5F-MMB-PICA, also increased the efficacy, which was comparable to 5F-NNEI. Most strikingly, adding an extra methyl group at the 3 position of 5F-MMB-PICA, resulting in 5F-MDMB-PICA, dramatically increased both the efficacy and potency of the compound. For comparison, we also included concentration response curves for additional structurally unrelated cannabinoid agonists (see Supplementary Figure 2, Table 1). Among them, Δ^9^-THC, anandamide, and 2-arachidonoylglycerol showed a lower potency and efficacy than CP55940.

### SAR for β-arrestin 2 recruitment by 5F-pentylindoles is similar to that for G protein engagement

β-arrestins play a crucial role in both desensitizing receptors and initiating their own signaling cascades [44]. Moreover, β-arrestin 2 is also involved in CB1R desensitization and internalization [45–49]. Therefore, we compared G protein engagement and β-arrestin recruitment by CB1Rs using 5F-pentylindole agonists. To ensure an unbiased comparison in a heterologous system, the same CB1R Rluc-fusion construct used for the CB1R-Gi1 coupling was also used for β-arrestin 2 recruitment (see Methods).

It has been reported that agonists have lower potencies for β-arrestin recruitment compared to G protein coupling [50]. As expected, most of the agonists tested in this study demonstrated ∼3-fold weaker potencies for β-arrestin 2 recruitment than for G_i1_ engagement (Figure 2, Table 1). Nonetheless, the potency ranking among SCRAs for β-arrestin 2 recruitment was similar to that observed for G_i1_ engagement. Specifically, the ketone-to-amide linker change resulted in decreased potency, as observed in the comparison between AM2201 and 5F-NNEI. Interestingly, among all the tested 5F-pentylindoles with amide-linker, only 5F-MDMB-PICA showed a higher potency than AM2201. Furthermore, similar to the G_i1_ engagement assay, the addition of one extra methyl group from 5F-MMB-PICA to 5F-MDMB-PICA led to a more than 6-fold increase in potency. The 5F-SDB-006 ligand had the lowest efficacy among the tested 5F-pentylindoles, while 5F-MDMB-PICA showed a much higher efficacy than 5F-MMB-PICA. Additionally, dose response profiles for other structurally unrelated cannabinoid agonists are displayed for comparison (Supplementary Figure 3, Table 1). Among them, THC, anandamide, and 2-arachidonoylglycerol showed lower potency and efficacy than CP55940.

**Figure 2.**
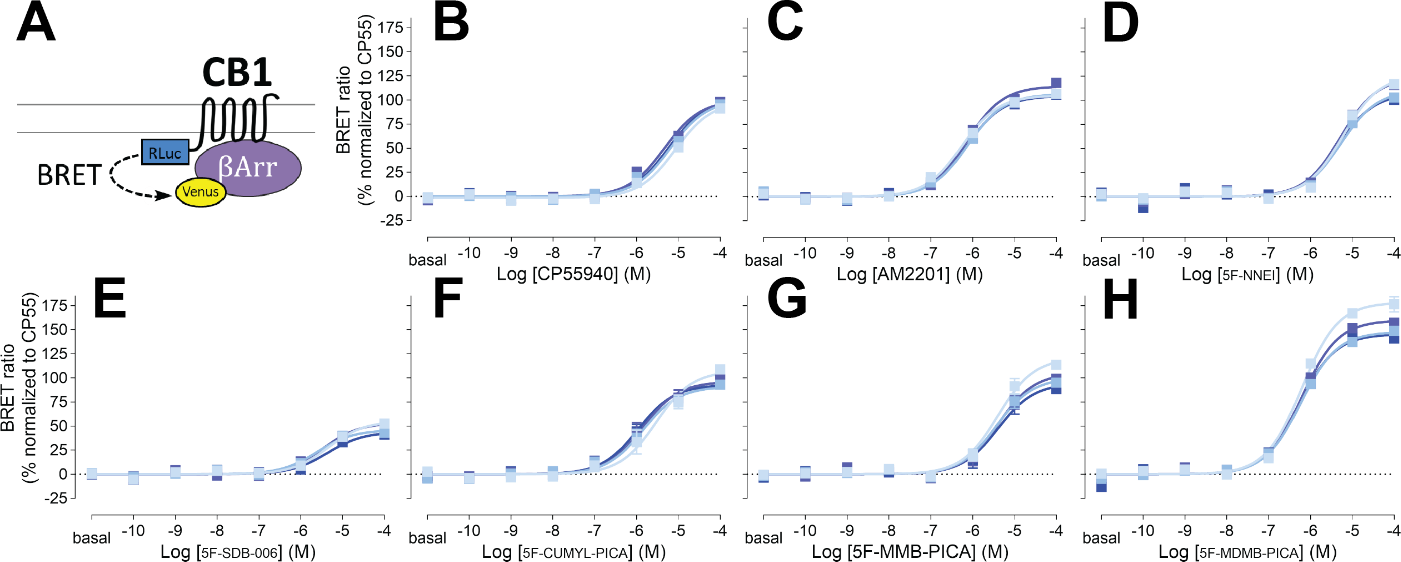
β-arrestin 2 recruitment BRET. Drug-induced recruitment BRET between CB1R-Rluc and β-arrestin 2-Venus (A) measured in response to CP55940 (B), AM2201 (C), 5F-NNEI (D), 5F-SDB-006 (E), 5F-CUMYL-PICA (F), 5F-MMB-PICA (G), 5F-MDMB-PICA (H) at various time points (2, 16, 30, 44 min light to dark blue). Concentration-response curves are plotted as a percentage of maximal response by CP55940 at each time point and presented as means ± SEM of *n* ≥ 3 independent experiments.

### No significant bias was observed between G protein engagement and β-arrestin recruitment

Signaling bias towards G protein activation or β-arrestin recruitment can result in downstream responses that deviate from those of balanced endogenous agonists. These differences could underlie the unique physiological outcomes reported for the use of SCRA in humans. We plotted the efficacies and potencies of various SCRAs and reference agonists for G_i1_ engagement and β-arrestin 2 recruitment (Figure 3). Many of the 5F-pentylindoles showed similar profiles for both signaling pathways, including 5F-MMB-PICA and 5F-MDMB-PICA (orange and green symbols in Figure 3). Of these, 5F-MDMB-PICA showed the highest efficacy and potency for both pathways. CP55940 noticeably showed a lower potency for β-arrestin 2 recruitment relative to other compounds (blue symbol, Figure 3). None of the agonists showed significant bias towards either pathway (i.e., with a bias factor >2.00 or <-2.00; Table 2).

**Figure 3.**
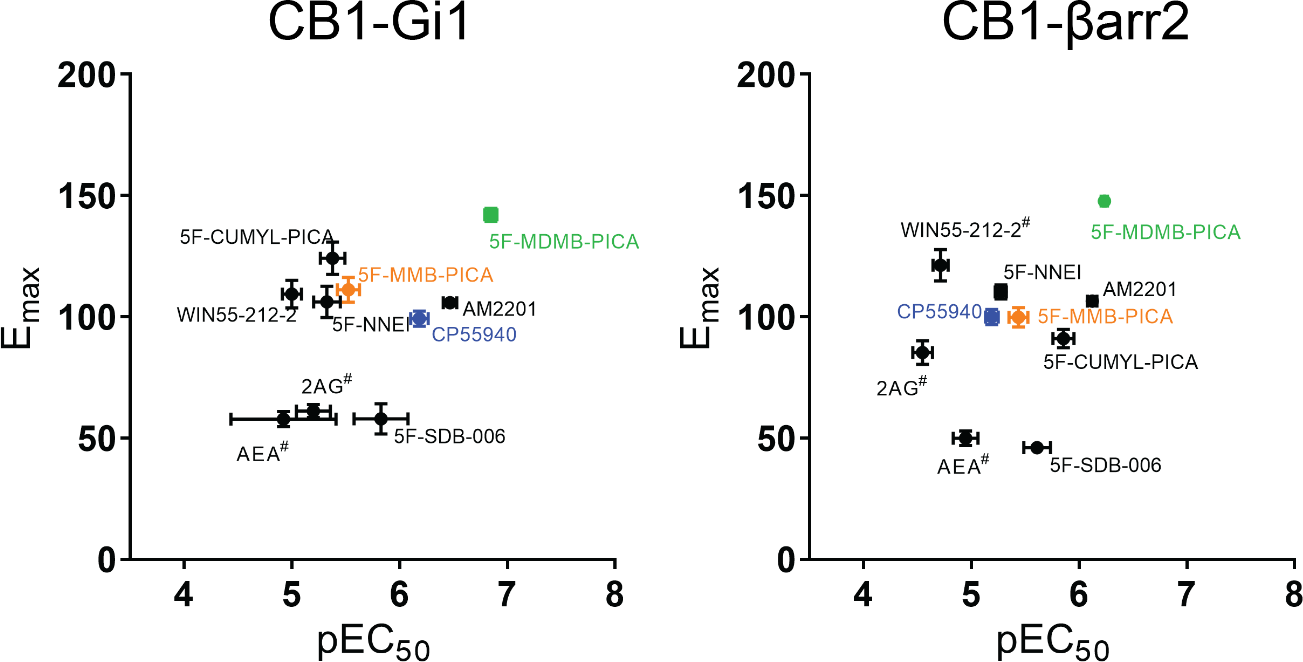
E_max_-pEC_50_ plot. E_max_-pEC_50_ comparison between CB1R-Gi1 engagement and β-arrestin 2 recruitment ^#^ Y-axis cut-off value at the highest concentration is shown for its E_max_.

**Table 2.**
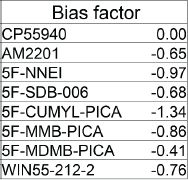
Coupling bias between Gi1 and β-arrestin 2 among the tested CB1R agonists.

In both the G_i1_ engagement and β-arrestin 2 recruitment assays, 5F-MDMB-PICA demonstrated the highest efficacy among all the compounds tested. This represented an increase of more than 40% compared to CP55940 and 5F-MMB-PICA, which has only one less methyl group in the head moiety. Therefore, to further characterize the actions of SCRAs at the CB1R, we focused on comparing 5F-MDMB-PICA and 5F-MMB-PICA in subsequent *ex vivo* recording and molecular simulation studies.

### Significant efficacy difference was observed between 5F-MDMB-PICA and 5F-MMB-PICA at presynaptic CB1Rs in the hippocampus

CB1Rs are expressed at the presynaptic terminals of neurons to regulate neurotransmitter release in the brain. In the hippocampus, CB1Rs are found on pyramidal neuron axonal terminals, known as Schaffer collateral/commissural (Scc) fibers projecting from the CA3 to CA1 regions [51]. Activation of CB1Rs results in Gi/o protein-mediated inhibition of voltage-dependent Ca^2+^ channels, leading to a reduction in glutamate release from Scc fibers in CA1 [51–53]. We utilized extracellular recordings of the rising slopes of field excitatory postsynaptic potentials (fEPSPs), evoked by electrical stimulation of Scc fibers in hippocampal brain slices, to compare the effects of 5F-MDMB-PICA and 5F-MMB-PICA on glutamate release [54].

As previously demonstrated with other synthetic CB1R agonists [55], fEPSPs were significantly reduced by bath application of either 5F-MDMB-PICA or 5F-MMB-PICA (1 µM, Figure 4A). As depicted in a time-course plot of fEPSP slope, the effects of both agonists are apparent within 10 min of application with calculated tau values of 6.34 and 8.80 min for 5F-MDMB-PICA and 5F-MMB-PICA, respectively (Figure 4B). However, the maximal inhibition produced by 5F-MDMB-PICA was significantly greater than that observed with 5F-MMB-PICA. Thus, concentration-response curves revealed significantly different E_max_ values for the two SCRAs (Figure 4C, t-test *P = 0.0255*). In order to confirm the CB1R-dependence of these effects, we applied the CB1R-selective antagonist AM251 after the effects of the agonists were stable. We found that AM251 fully reversed the effects of both agonists (Supplementary Figure 4).

**Figure 4.**
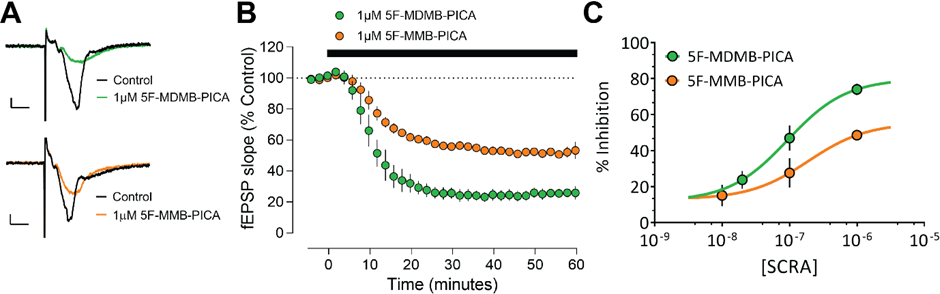
Brain slice electrophysiology. Concentration-dependent inhibition of hippocampal glutamate release by 5F-MDMB-PICA and 5F-MMB-PICA. (A) Representative averaged traces for 5F-MDMB-PICA (upper) and 5F-MMB-PICA (lower), (B) Summary time course of recordings (*n* ≥ 3 recordings) demonstrating the effect of SCRAs. (C) Concentration–response curve for 5F-MDMB-PICA and 5F-MMB-PICA (*n* ≥ 3 slices per concentration). The pEC_50_ was calculated to be 7.20 and 6.72 for 5F-MDMB-PICA and 5F-MMB-PICA respectively. Data points are presented as means ± SEM.

### Adding an extra methyl group to the ligand head moiety changes the dynamics of the agonist binding pocket

The *in vitro* and *ex vivo* assays demonstrated that 5F-MDMB-PICA is significantly more efficacious than 5F-MMB-PICA, with the latter’s efficacy being comparable to that of the reference full agonist CP55940 (Figure 3). This observation led us to investigate the molecular mechanism underlying 5F-MDMB-PICA’s higher efficacy at the CB1R. That only the addition of a single methyl group to the head moiety of 5F-MMB-PICA could cause such a large increase in 5F-MDMB-PICA’s efficacy permitted us to investigate how this affects ligand-receptor interactions, and the propagation to receptor/G-protein coupling. To do this, we conducted comparative molecular modeling and simulations of the CB1R bound to each of these SCRAs.

We used the CB1R/Gαi protein complex bound with MDMB-FUBINACA (Figure 5A; PDB 6N4B), as the template to construct the CB1R/5F-MDMB-PICA and CB1R/5F-MMB-PICA complex models (see Methods). In the structure of 6N4B, the head moiety and linker of MDMB-FUBINACA occupy a pocket that is enclosed by residues from transmembrane helices (TMs) 1, 2, and 7, while its *p*-fluorobenzyl tail is enclosed by TMs 3, 5, 6, and second extracellular loop (ECL2). Notably, 5F-MDMB-PICA and MDMB-FUBINACA share the same head moiety, while 5F-MMB-PICA has one less methyl group than these two ligands (Supplementary Figure 5). Therefore, we assumed that 5F-MDMB-PICA or 5F-MMB-PICA would have a similar head-tail orientation as MDMB-FUBINACA in the ligand binding pocket (Supplementary Figure 5), and therefore selected their corresponding docking poses accordingly to construct the complex models. As a control, we also simulated the CB1R/MDMB-FUBINACA model (Table 3). During our simulations, 5F-MMB-PICA and 5F-MDMB-PICA maintained their binding poses in the same orientation as MDMB-FUBINACA (Figure 5C) but exhibited some variations in both the local interactions within the binding pocket and the resulting global receptor conformation.

**Figure 5.**
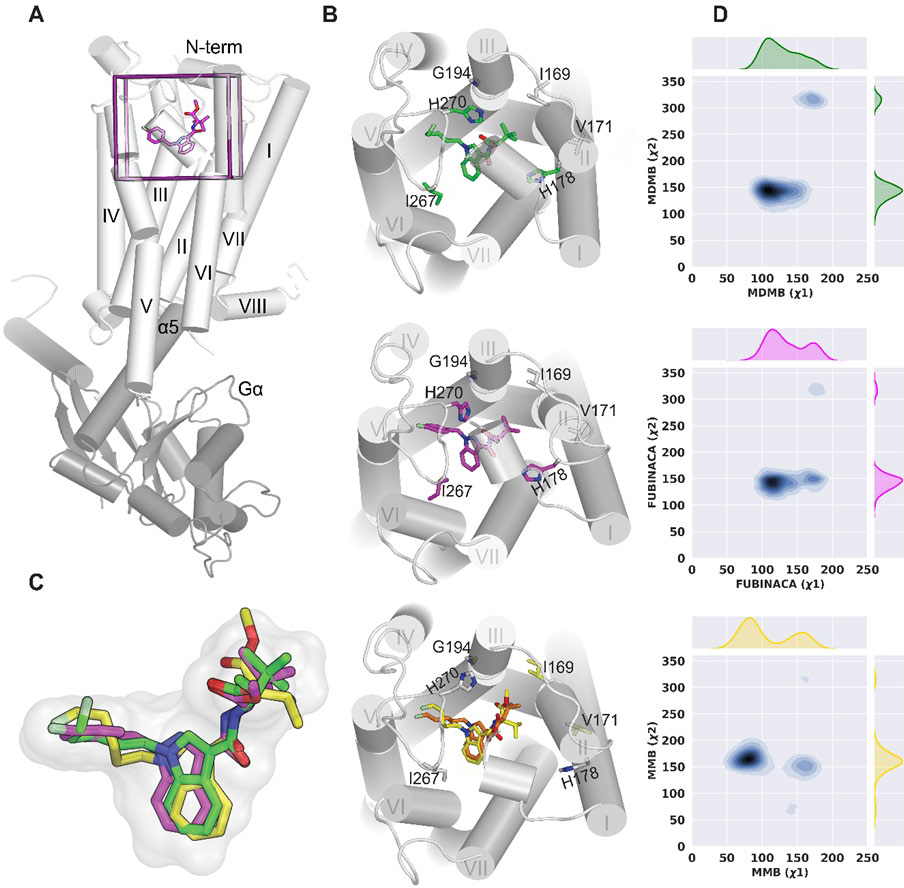
Ligands’ dihedral angle and binding pocket. (A) The zoom-out view of the receptor (grey) bound with Gαi (dark grey) and MDMB-FUBINACA (magenta). (B) The binding mode of 5F-MDMB-PICA (green), MDMB-FUBINACA (magenta), and 5F-MMB-PICA (yellow and orange) and their associated binding site residues that show differences in contact frequency between 5F-MDMB-PICA, MDMB-FUBINACA, and 5F-MMB-PICA. The residues that show differences are color coded correspondingly. (C) The zoom-in view of the superimposed ligands in the binding site. They occupy the same region in binding pocket but 5F-MMB-PICA picks two distinct conformers for head moiety. (D) The 2D map of head moiety dihedral angles (Supplementary Figure 1).

**Table 3.** Summary of simulated conditions and simulation lengths.

| Ligand | Number of trajectories | Length of trajectories ($\mu s$) |
| --- | --- | --- |
| 5F-MDMB-PICA | 5 | 14.5 |
| 5F-MMB-PICA | 6 | 14.5 |
| MDMB-FUBINACA | 3 | 6.6 |

Based on the equilibrated simulation results, we first assessed the impact of the extra methyl group on the ligand binding poses and measured two dihedral angles in the head moiety for each bound ligand (designated as *x*1 and *x*2, Supplementary Figure 5). The distribution of *x*1 revealed that 5F-MMB-PICA had two distinct modes in the binding site, with *x*1 (∼50 ≤ peak 1 ≤ ∼115 and ∼135 ≤ peak 2 ≤ ∼180), resulting from the rotation of the head moiety around the N-C bond (Figure 5D). In contrast, there appeared to be minimal energy barriers for the head moieties of the bound 5F-MDMB-PICA and MDMB-FUBINACA to transition between these two modes. The *x*2 dihedral angle did not exhibit any noticeable differences among the three ligands.

We hypothesized that the distinct dynamics of the ligand head moiety in the binding pocket might impact their interactions with the receptor. Consequently, to evaluate the effects of the extra methyl group on the dynamics of the binding site, we calculated the contact frequency and residues’ sidechain dihedral angles for each ligand (see Methods). The outcomes revealed that contact residues were present in TM2, TM3, ECL2, TM5, TM6, and TM7. TM2 and TM3 had a more significant role in ligand binding interactions than other segments (Supplementary Table 1). Notably, our studies revealed clear distinctions among the three simulated conditions investigated. The contact frequency and sidechain dihedral angle analyses suggested that 5F-MDMB-PICA and MDMB-FUBINACA, with the same head moiety, shared similarities for most of their contacting residues, and their differences with 5F-MMB-PICA were similar (Supplementary Table 1). Specifically, several residues from TM2 (I169^2.56^, V171^2.58^, S173^2.60^, and H178^2.65^), TM3 (G194^3.30^), and ECL2 (I267^ECL2^, P269^ECL2^, and H270^ECL2^), which all interact with the ligand head moieties, showed similar differences.

In particular, our analysis shows that H178^2.65^ plays a crucial role in ligand interactions, with both 5F-MDMB-PICA and MDMB-FUBINACA interacting more frequently (∼100%) with this residue than 5F-MMB-PICA (46%). In addition, our results reveal that 5F-MDMB-PICA interacts more with I267^ECL2^, I269^ECL2^, and H270^ECL2^ compared to 5F-MMB-PICA, as do I267^ECL2^ and H270^ECL2^ in the case of MDMB-FUBINACA (Supplementary Table 1, Figure 5B). Furthermore, we observed that the dynamics of the ligands in the binding pocket can induce changes in the sidechain conformations of certain residues, as highlighted in Supplementary Table 1. For example, the *X*1 rotamer of S173^2.60^ for 5F-MDMB-PICA and MDMB-FUBINCA has more than 70° difference from that 5F-MMB-PICA. Additionally, there are noticeable differences in the sidechain conformation of P269^ECL2^ between 5F-MDMB-PICA/5F-MMB-PICA and MDMB-FUBINACA/5F-MMB-PICA, (Supplementary Table 1) which could be related to their different puckering states.

### The additional methyl group in head moiety causes conformational changes in TM1 and TM2

To investigate the conformational changes beyond the ligand pocket, we first compared the available CB1R structures in the active and inactive states. The CB1R shares activation features with other class A GPCRs, including outward movement of the intracellular portion of TM6 (TM6i) and inward movement of TM7i, accompanied by changes in conserved motifs [56]. Our quantitative analysis, shown in Supplementary Figure 6, demonstrates these movements between the active CB1R-Gαi structures (PDB: 6N4B, 6KPG, and 7WV9) [28–30] and the inactive structures (PDB: 5U09 [31] and PDB: 5TGZ [32]). In addition to these features, CB1R also exhibits inward movements of the extracellular portion of TM1 (TM1e) and TM2e with the displacement of the N terminus upon activation [28].TM2 plays a distinct role in CB1R activation by engaging the ligand in hydrophobic/polar interactions with TM2e and stabilizing the agonist binding conformation[28, 32]. However, 6N4B shows another interesting feature at the extracellular domain which is the inward movement of TM6e towards TM1e and TM2e (Supplementary Figure 6).

For our simulation results, after performing RMSD-based clustering of CB1R TM domain and conducting the PIA-GPCR analysis, we observed conformational differences in the TM1e and TM2e region between CB1R/5F-MDMB-PICA and CB1R/5F-MMB-PICA (Figure 6A). Specifically, we noted that in CB1R/5F-MDMB-PICA, TM1e and TM2e moved inward towards the center of the ligand binding pocket, in contrast to their configuration in CB1R/5F-MMB-PICA. As a result of the inward movement of TM2 in CB1R/5F-MDMB-PICA, key residues in TM2e, such as H178^2.65^, had more frequent interactions with the head moiety of 5F-MDMB-PICA in the binding pocket (Supplementary Table 1), thus stabilizing the bound 5F-MDMB-PICA.

**Figure 6.**
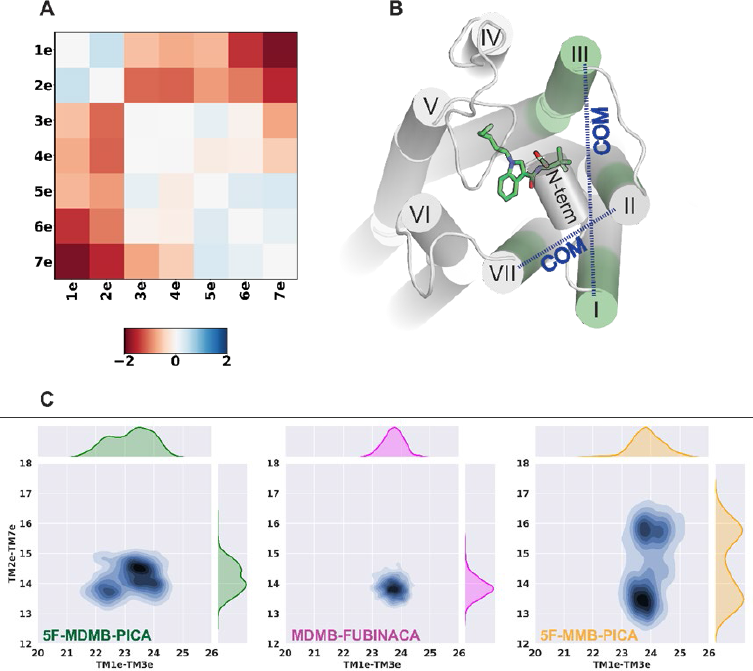
PIA-GPCR analysis and TMs’ COM distribution. (A) PIA-GPCR COM results for TMs’ extracellular domain (ΔTMe(COM) = TMe(COM)_5F-MDMB-PICA_ – TMe(COM)_5F MMB-PICA_). (B) The 5F-MDMB-PICA in the binding pocket. The highlighted transmembrane sub-segments are extracellular part of TM1, TM2, TM3, and TM7 (denoted as TM1e, TM2e, TM3e, and TM7e). The blue dashed arrows show the COM distance between TM1e – TM3e and TM2e – TM7e. (C) The 2D distribution of the COM distances. 5F-MMB-PICA shows two distinct distributions for TM2e – TM7e COM distance which seems to be the result of the changes in head moiety seen for this ligand.

To further characterize the conformational changes in this receptor region that accommodate the head groups of the SCRAs, we analyzed the center-of-mass (COM) distances between two pairs of transmembrane subsegments: TM1e – TM3e and TM2e – TM7e (Figure 6B). The COM distance distributions between TM2e – TM7e shows two distinct peaks for 5F-MMB-PICA compared to 5F-MDMB-PICA and MDMB-FUBINCA (Figure 6C). This could be due to the two conformations that the head moiety of 5F-MMB-PICA can adopt. The extra methyl group of 5F-MDMB-PICA results in a more stable conformation for its head moiety, which is associated with the changes in the dynamics of the extracellular regions of the receptor surrounding the head moiety.

### Divergence in the allosteric interaction network between 5F-MDMB-PICA and 5F-MMB-PICA

We then carried out a correlation-based network analysis (see Methods) to compare how the bindings of 5F-MDMB-PICA and 5F-MMB-PICA may induce different conformational changes from the ligand binding pocket to the receptor-G protein interface. Our results identified several significant differences between CB1R/5F-MDMB-PICA and CB1R/5F-MMB-PICA conditions, as illustrated in Figure 7A.

**Figure 7.**
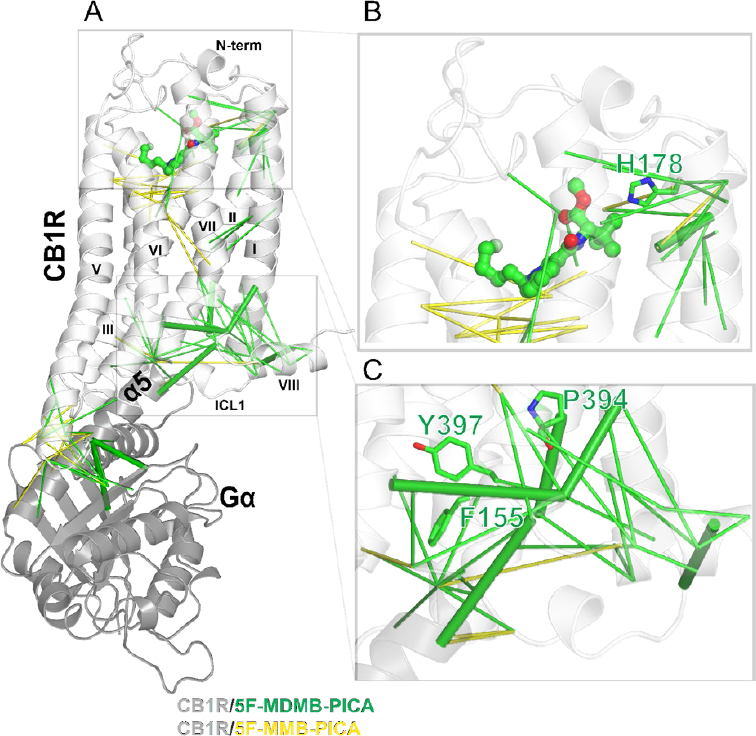
Correlation-based network pathway. (A) The CB1R structure bound with 5F-MDMB-PICA with final correlated pairs mapped on the structure. The green pairs are representative pairs for 5F-MDMB-PICA and the yellow pairs are representative for 5F-MMB-PICA. There is a noticeable difference between these two ligands signaling pathway. The CB1R/5F-MDMB-PICA network includes a pathway consisting of eight strongly correlated pairs (solid green lines) which connect the E133^1.49^-L399^7.55^ (TM1i – TM7i) changes to the changes at the extracellular-middle domain, F174^2.61^-D176^2.63^ (TM2m – TM2e), the intracellular domain, A160^2.47^-A398^7.54^ (TM2i – TM7i), H143^1.59-^F408^H8^ (TM1i – H8), T344^6.36^-L399^7.55^ (TM6i – TM7i), H406^H8^-R409^H8^ (H8 – H8), and the changes between the intracellular domain and Gα,T313^ICL3^-S265<u>^Gα^</u> (ICL3 – Gα), T313 ^ICL3^-E318^Gα^ (ICL3 – Gα), and R400^7.56^-K349^Gα^ (TM7i – Gα). E133^1.49^-L399^7.55^ (TM1i – TM7i) is at the core of these correlated pairs. There is no such pathway for CB1R/5F-MMB-PICA. (B) The TM2e key residue H178^2.65^ contributes to the 5F-MDMB-PICA signaling pathway. (C) The TM2i residue (F155^2.42^) and TM7i NPxxY residues (P394^7.50^ and Y397^7.53)^ are unique to 5F-MDMB-PICA network pathway

Specifically, TMs 1, 2, and 7 are more engaged in the allosteric network for CB1R/5F-MDMB-PICA, while TMs 3,4, and 5 are more involved for the CB1R/5F-MMB-PICA. The analysis shows that ∼45% of TM2 residues are part of the strongly correlated network for CB1R/5F-MDMB-PICA while only 14% of TM2 residues are involved in the network for CB1R/5F-MMB-PICA. Notably, the two key residues in CB1R binding (H178^2.65^) (Figure 7B) and activation (F155^2.42^) (Figure 7C) [29] contribute to the network for CB1R/5F-MDMB-PICA but not for CB1R/5F-MMB-PICA. Similarly, for TM7, the contribution of residues for CB1R/5F-MDMB-PICA and CB1R/5F-MMB-PICA is 44% and 15%, respectively. Among these residues, P394^7.50^ and Y397^7.53^ from the NPxxY motif are unique to the CB1R/5F-MDMB-PICA network (Figure 7C). As reviewed and described above, TM2 and TM7 have been shown to play critical roles in CB1R activation. Therefore, such distinct TM and residue involvements between these two conditions may be associated with the different efficacies of 5F-MDMB-PICA and 5F-MMB-PICA at CB1R.

Additionally, the number of correlated residue pairs for CB1R/5F-MDMB-PICA is almost twice that of CB1R/5F-MMB-PICA. The results showed pairs with strong correlations that connect the changes in binding pocket between the 5F-MDMB-PICA and receptor residues to other pairs (receptor-receptor, receptor-Gαi). While these residue pairs do not form a complete pathway for either CB1R/5F-MDMB-PICA or CB1R/5F-MMB-PICA, the rigid body movement of the receptor regions between them likely propagate the impact.

Thus, our results underscore the contributions of the highly conserved NpxxY motif [56], as well as the key residues H^2.65^ and F^2.42^ [29] in the allosteric network that may be associated with receptor activation.

## DISCUSSION

Here we demonstrated substantial increase in the efficacy and potency of a 5-fluoropentylindole SCRA through the addition of a single methyl group to the head moiety (i.e., 5F-MDMB-PICA). This modification led to superagonism, which refers to the phenomenon where agonist efficacy surpasses that of endogenous agonists [33]. Although “superagonism” has a clear definition, its interpretation can often be complicated when applied to assays that not directly detect receptor activation status. These results can be confounded by signal amplification when large receptor reserves are present, which can lead to inaccurate representation of the intrinsic efficacy of the agonist [57]. The BRET assays utilized in this study rely on the direct coupling between a receptor and a transducer, which can effectively avoid signal amplification and detect transducer engagement or recruitment resulting from receptor activation. However, compared to the cAMP assay, the use of fusion constructs in BRET assays, which enable specificity and direct coupling, may lead to comparatively lower potencies. Nonetheless, a comparison between the test ligands and a reference CP55940 ensures proper calibration of the relative potencies of SCRAs. Our finding of 5F-MDMB-PICA as a superagonist at the CB1R is supported not only by our own electrophysiological results but also by previous studies in cellular functional assays measuring membrane potential [8, 40], and the cannabinoid triad behavioral assays [58]. Notably, the presynaptic inhibition measured using field recordings in brain slices is known to be sensitive to differences in pharmacological properties of CB1R ligands that possess an adequate “ceiling” to characterize high efficacy agonists.

Addition of a methyl substituent, although not a dramatic structural change, is known to yield profound ligand-receptor interactions in many cases [59, 60], as observed here with the methyl-3-methylbutanoate to methyl-3,3-dimethylbutanoate modification (i.e., MMB- to MDMB-change). Extensive interactions between the methyl-3,3-dimethylbutanoate head moiety and the TM2e region were demonstrated in the cryo-EM structure [28], and our computational modeling demonstrated how these interactions can significantly affect the potency and efficacy of 5F-MDMB-PICA compared to 5F-MMB-PICA. Our simulation results also indicated additional interactions of 5F-MDMB-PICA with ECL2 residues, compared to 5F-MMB-PICA, which maybe also relate to its higher potency and efficacy. This finding aligns with previous mutagenesis work of the CB1R, which have highlighted the critical role of ECL2 in ligand binding [28, 61], G protein coupling, and receptor trafficking [61]. In particular, alanine scanning mutations on both N- and C-terminal regions of ECL2 have consistently resulted in more than a 10-fold increase in EC_50_ with significantly reduced E_max_ values, suggesting the indispensability of these residues for both ligand binding and receptor activation [61].

Since the initial reports of illicit usage around 2010, there has been an increasing recognition of the association between the potency and efficacy of SCRAs at CB1Rs and their clinically adverse effects [62]. Several studies have focused on assessing the pharmacological properties of the head, linker, core, and tail moieties comprising SCRAs [63, 64]. Consistent with the findings of our current study, previous data have also indicated that even minor structural variations within SCRAs can lead to significant efficacy changes, thereby potentially influencing clinical toxicity [6, 25, 65]. Among them, 5F-MDMB-PICA, a schedule-I drug, has been identified as one of the most prevalent SCRAs in United States, as of 2019 [66]. Fatal events have been associated with the use of this drug [67, 68], as well as reports of notably strong depressant effects [69], and other toxic interactions [70]. Our present findings suggest that even minor structural modifications in newly emerging SCRAs, particularly in relation to the presence of substituted head moieties, can strongly increase SCRA efficacy, and potentially contribute to the adverse effects occurring with human consumption of these illicit compounds. Our study can be used to further our understanding of how specific alterations in cannabinoid receptor ligands may contribute to their toxicity in humans and the potential life threatening consequences of their use.

## METHODS AND MATERIALS

### Cell culture, constructs, BRET, analysis

Variations of bioluminescence resonance energy transfer (BRET) assay were performed to detect ligand-induced receptor-signaling protein coupling events. A constant amount of plasmid cDNA (15 μg) was transfected into human embryonic kidney cells 293 T (HEK-293T) using polyethylenimine (PEI; Sigma) in a 1:2 weight ratio in 10 cm plates. Cells were maintained in culture with Dulbecco’s modified Eagle’s medium (DMEM) supplemented with 10% fetal bovine serum (FBS, Atlanta), 2 mM L-glutamine (Gibco), and 1% penicillin streptomycin (Gibco) and kept in an incubator at 37 °C and 5% CO_2_. The transfected amount and ratio among the receptor and heterotrimeric G proteins were tested for the optimized dynamic range in drug-induced BRET. Experiments were performed approximately 48 h post-transfection. As reported previously [71], cells were collected, washed, and resuspended in phosphate-buffered saline (PBS). Approximately 200,000 cells/well were distributed in 96-well plates, and 5 μM coelenterazine H (luciferase substrate) was added to each well. One minute after the addition of coelenterazine, CB1R ligands were added to each well. Two different configurations of BRET were used: (i) Gαi1-engagement and (ii) β-arrestin-2 recruitment. (i) G_i1_ protein engagement assay uses Rluc-fused receptor and mVenus-fused Gαi1 for a resonance energy transfer (RET) pair. Untagged Gβ1 and Gγ2constructs were cotransfected. (ii) β-arrestin-2 recruitment uses D2R-Rluc-β-arrestin-2-Venus for a RET pair. GRK2 was cotransfected to assist an enhanced phosphorylation required for the β-arrestin-2 recruitment. The donor luminescence as well as the acceptor fluorescence were always quantified for consistent expression levels across different experiments such that no significant expression differences. For fluorescence measurement, Venus was excited at 500 nm and measured at an emission wavelength of 530 nm over 1 s, using a Pherastar FSX plate reader (BMG Labtech, Cary, NC, USA). For kinetic experiments, cells were incubated at 37 °C within the Pherastar FSX plate reader (BMG Labtech, Cary, NC, USA) with BRET signal measurements taken at various time-points ranging from 2–46 min. The BRET1 signal from the same batch of cells was calculated as the ratio of the light emitted by Venus (530 nm) over that emitted by coelenterazine H (485 nm). BRET change was defined as BRET ratio for the corresponding drug minus BRET ratio in the absence of the drug. *E*_max_ values are expressed as the basal subtracted BRET change and in the dose– response graphs. Data and statistical analysis were performed with Prism 9 (GraphPad Software). Bias factors between Gi1-engagement and β-arrestin-2 were calculated as previously described [71, 72].

### Brain Slice Electrophysiology

Experiments were performed as described previously, with additional modifications [55, 71]. Brain slices were prepared from adult wildtype C57BL/6J mice (male and female) that were anesthetized with isoflurane and euthanized by decapitation. Prior to decapitation, mice were intracardially perfused with modified artificial cerebral spinal fluid (mACSF) containing (in mM): 92 NMDG, 20 HEPES, 25 glucose, 30 NaHCO_3_, 1.2 NaH_2_PO_4_, 2.5 KCl, 5 sodium ascorbate, 3 sodium pyruvate, 2 thiourea, 10 MgSO_4_, 0.5 CaCl_2_, 300–310 mOsm, at pH 7.3–7.4. The brains were sectioned in cold mACSF, saturated with 95% O_2_ and 5% CO_2_ (carbogen), and transverse hippocampal slices (220 µm) were obtained using a vibrating tissue slicer. The slices were immediately placed in the same buffer and maintained at 32 °C for 10 min, before moving them to a holding chamber filled with carbogen saturated ACSF (holding ACSF) containing, in mM: 92 NaCl, 20 HEPES, 25 glucose, 30 NaHCO_3_, 1.2 NaH_2_PO_4_, 2.5 KCl, 5 sodium ascorbate, 3 sodium pyruvate, 2 thiourea, 1 MgSO_4_, 2 CaCl_2_, 300–310 mOsm, at pH 7.3–7.4. During electrophysiological recordings, slices were continuously perfused at 2 mL/min with carbogen-saturated ACSF containing (in mM): 124 NaCl, 2.5 KCl, 1.25 NaH_2_PO_4_, 1 MgCl_2_, 26 NaHCO_3_, 11 glucose, 2.4 CaCl_2_, 300–310 mOsm, at pH 7.3–7.4. The temperature of the recording chamber was maintained at 31–32 °C. Brain slices were visualized using an upright microscope to confirm placement of an extracellular recording electrodes (3–5 MΩ), backfilled with ACSF, within the *stratum radiatum* of area CA1. A bipolar stimulating wire was positioned in area CA3A to activate Scc fibers and evoke fEPSPs. Extracellular recordings were made using a MultiClamp 700B amplifier (10 kHz low-pass Bessel filter) and Digidata 1550B (20 kHz digitization) with pClamp 11 software (Molecular Devices). As we have previously found that endogenous adenosine can disrupt CB1R-mediated inhibition of glutamate release in the hippocampus [71], we included the adenosine A_1_ receptor antagonist, caffeine (50 μM) in the ACSF throughout incubation and recordings.

The following numbers of slices were recorded for the drug perfusion conditions listed: 2×10^8^, 10^−7^, 10^−6^ M 5F-MDMB-PICA (6 slice recordings from 4 animals); 10^−8^, 10^−7^, 10^−6^ M 5F-MMB-PICA (5 slice recordings from 4 animals); 2×10^−8^ M 5F-MDMB-PICA + 10^−6^ M AM251 (5 slice recordings from 5 animals); 10^−6^ M 5F-MMB-PICA + 3×10^−6^ M AM251 (6 slice recordings from 6 animals). All data are reported as mean ± SEM. Data were analyzed in Clampex and statistically compared with Prism 9 (GraphPad Software) by paired t-test.

### Molecular dynamics simulations

We used the (MDMB-FUBINACA)-CB1R-Gαi crystal structure (PDB 6N4B) [28] as the starting point for our modeling and molecular dynamics (MD) simulations. The CB1R missing residues on the N-terminus (residues 104-108), extracellular loop 2 (ECL2) (residues 258-263), and helix-8 (H8) (residues 412-414) were added using MODELER (version 9.24) with CB1R structure (PDB 5XRA) [27] as the template.

The structure was further processed and refined using the Protein Preparation Wizard and Ligand Preparation implemented in Maestro (Schrödinger suite 2019-4). In addition, the residues D163^2.50^ and D213^3.49^ were protonated to their neutral forms as assumed in the active state of rhodopsin-like GPCRs [73–75].

Initial poses of 5F-MDMB-PICA and 5F-MMB-PICA were selected by analyzing the results of docking the ligands into the prepared (MDMB-FUBINACA)-CB1R binding site using the Induced-Fit Docking (IFD) protocol [76] (Schrödinger suite 2019-4). Using Desmond System Builder (Schrödinger suite 2019-4), CB1R models were placed into explicit 1-palmitoyl-2-oleoyl-sn-glycero-3-phosphocholine lipid bilayer (POPC) with the standard membrane-protein orientation provided by the Orientation of Proteins in Membranes (OPM) database [77]. Simple point charge (SPC) water model [78] was used to solvate the system, charges were neutralized, and 0.15 M NaCl was added. The total system size was ∼177000 atoms. The OPLS3e force field (PMID: 30768902) was used throughout this study. The initial parameters for 5F-MDMB-PICA and MDMB-FUBINACA based on the default atom typing of OPLS3e was further optimized by the force field builder (Schrödinger suite 2019-4). Desmond MD system (D. E. Shaw Research, New York, NY) [79] was used for the unbiased all-atom MD simulations.

We applied similar simulation protocols for GPCRs [80] to minimize and equilibrate the system. The initial energy minimization was followed by equilibration with the restraints on the ligand heavy atoms and protein backbone atoms. In the production runs, we used the NPT ensemble with constant temperature at 310 K maintained with Langevin dynamics and 1 atm constant pressure was achieved by hybrid Nosé-Hoover Langevin dynamics on an anisotropic flexible periodic cell with a constant surface tension (x-y plane). In production runs, all restraints on the CB1R were released. However, restraints on **α**5 and N-terminus segments of Gα were kept for 1.2 *μ*s and then released.

### Clustering analysis

We performed root-mean-square deviation (RMSD)-based clustering of our aggregated MD simulation trajectories to study the conformational dynamics of protein domains. The clustering is based on all-to-all RMSD of the backbone of transmembrane segments (TMs). The computed all-to-all RMSD matrix was then subjected to the K-means clustering algorithm [81]. For each ligand, we combined frames collected from all independent trajectories, then sampled 10000 frames with replacement. Finally, we merged the sampled frames from each ligand and used them to calculate the all-to-all RMSD.

To estimate the number of clusters, we used the Gap Statistic method (1) [81]. In this technique the change in within-cluster sum of squares error is compared with the expected value under an appropriate null distribution.

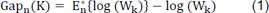

where 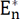 is the expectation of log (W_k_) and n is the size of the sample from the reference distribution. 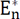 {log (W_k_)}can be estimated as follows: 1) Uniformly generate the reference distribution 2) Draw B number of Monte Carlo samples from the reference distribution 3) Determine the log (W_k_) for each sample 4) Estimate 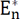{log (W_k_)} by an average of B copies log (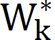). We can then calculate the gap statistic by following two steps: 1), change the number of clusters form k = 1, 2, …, K, cluster the observed data, and for each k calculate the within-sum of squares of error (W_k_) 2) for each k, generate B reference data sets (b = 1, 2, …., B) from the uniform distribution and measure within-sum of squares of error 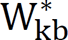. The gap statistic can be calculated as (2):

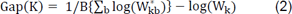

and using 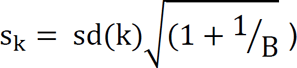, which is the simulation error in, the number of clusters is the smallest value for k such that Gap(K) ≥ Gap(K + 1) − s_k+1_

Frames from selected clusters (clustered frames*) were then mapped to the original frames and saved to perform further analyses (e.g., RMSD, dihedral angles, contact frequency, Protein Interaction Analyzer (PIA-GPCR) [82], and correlation-based network analysis).

Contact frequency: We considered the heavy atoms of binding site residues and each ligand to find the residues that were within 6 Å of each ligand to calculate the contact frequency for each ligand. If any of the ligands showed 50% or more frequency for any of the binding site residues, we tabulated the results for that residue for all studied ligands. We also calculated the differences between CB1R/5F-MDMB-PICA or CB1R/MDMB-FUBINACA, and CB1R/5F-MMB-PICA, respectively. The differences of 25% and more were highlighted.

PIA-GPCR: The following structural elements were defined for the analysis of coarse-grained interaction network of the hCB1. The “e”,” m”, and “i” symbols define the extracellular, middle, and intracellular section of TM region, respectively. TM1e (residues 113^1.29^-120^1.36)^, TM1m (residues 121^1.37^-129^1.45^), TM1i (residues 130^1.46-^144^1.60^), TM2e (residues 175^2.62^-179^2.66^), TM2m (residues 165^2.52^-174^2.61^), TM2i (residues 151^2.38^-164^2.51^), TM3e (residues 186^3.22^-195^3.31^), TM3m (residues 196^3.32-^202^3.38^), TM3i (residues 203^3.39^-220^3.56^), TM4e (residues 247^4.56^-253^4.62)^, TM4m (residues 241^4.50^-246^4.55^), TM4i (residues 229^4.38^-240^4.49^), TM5e (residues 272^5.36^-280^5.44^), TM5m (residues 281^5.45^-288^5.52^). TM5i (residues 289^5.53^-311^5.75^), TM6e (residues 363^6.55^-368^6.60^), TM6m (residues 355^6.47^-362^6.54^), TM6i (residues 335^6.27^-354^6.46)^, TM7e (residues 374^7.30^-382^7.38)^, TM7m (residues 383^7.39^-390^7.46^), TM7i (residues 391^7.47^-400^7.56^).

For PIA-GPCR sidechain rotamer analysis, the residues extracted from contact frequency analysis along with the residues reported in the experimental studies [27, 28] but not identified in contact frequency results were included.

RMSD, dihedral angles, and contact frequency were calculated with VMD (version 1.9.3), PIA-GPCR with our *in-house* scripts, and the Cα-Cα distances for network analysis with MDAnalysis (version 1.1.1).

### Correlation-based network analysis

We performed Pearson correlation analysis to identify the unique signaling pathways for ligands studied in our work. We divided the signaling pathway into three regions and for each region systematically scanned and included distance/dihedral measurements in our initial data set for network analysis based on the following considerations:

#### Ligand-binding site

Ligand polar groups play a significant role in ligand binding. In both 5F-MDMB-PICA and 5F-MMB-PICA the polar groups are distributed in head, linker, core and tail moieties. Three oxygen atoms in the head moiety, two nitrogen atoms in the linker and core, respectively, and one fluorine atom in the tail. Therefore, the dynamics of the ligand in the binding site can be characterized using the distances between these atoms and receptor residues. Clustered frames* were used to calculate the contact frequency between ligand and binding site residues. The residues within 6 Å of each polar atom that have 10% contact frequency were selected and the distances between those residues and the corresponding polar atom were included in network analysis. The dihedral angles of the ligands’ head moiety were also included in the analysis.

#### TMs site

Clustered frames* were used to measure the distances between the Cα-Cα of all 7TM residues. We then filtered out the Cα-Cα distances that their average value was larger than 12 Å and the rest were included in the final set of measurements for network analysis. The dihedral angles of the conserved motifs that play a significant role in GPCRs activation _(C6.47W6.48×6.49P6.50, D3.49R3.50Y3.51, P5.50I3.40F6.44, and N7.49P7.50×7.51×7.52Y7.53) were also measured_ and included in the analyses.

#### CB1R-Gαi site

Clustered frames* were used to systematically scan the distances between receptor residues (TMs + intracellular loops) and the Gα residues. The Cα-Cα distances that their average value was larger than 12 Å were filtered out and the rest were included in the final set of measurements.

The sample Pearson correlation coefficient (r) was calculated for all distance/dihedral pairs. We then applied three filtering criteria on Pearson correlation coefficient matrix

1. Threshold for r and only pairs with r ≥ 0.85 were kept.
2. Let’s show each correlated pair of distances as d_i_ = X_i_ - Y_i_ (distance i) and d_j_ = X_j_ - Y_j_ (distance j) where X and Y are the residue numbers for pairs i and j. If that pairs were kept.
3. For each set of correlated pairs that share one common distance, and the other distances include residues from the same region of TMs, only one pair with the maximum difference between residue numbers either |(X_i_ - X_j_)| or |(Y_i_ - Y_j_)| or both was kept.

The clustering and network analyses were repeated 10 times (bootstrap sampling with replacement). The final correlated pairs resulted from each bootstrap sampling which followed by clustering and network analysis were combined, and the common pairs were extracted for each ligand. The common pairs for each ligand were then mapped to the structure for visualization. These final pairs are considered as a hot region of allosteric pathway at the selected correlation threshold (0.85) for each ligand. To make sure the correlation between each pair is real (the change in variable m(t) (e.g., distance or dihedral angle) at time t_i_ and t_i+1_ is accompanied by the change in variable n(t) at the corresponding times), the time series for each final correlated pair were evaluated by plotting their distribution as a 2D map.

## Author Contributions

HY and LS designed the experiments and computations. HY, PT, AFH, and CL carried out the experiments, and performed the data analysis with LS, MHB, and CRL. RC carried out the computational modeling and simulations and performed the analysis with LS. All authors were involved in data interpretation. HY, RC, and LS prepared the initial manuscript, with contributions from all the authors in finalizing the manuscript.

## Acknowledgements

The work is supported by Brain & Behavior Research Foundation Young Investigator Grant (H.Y.) Support for this research was provided by the National Institute on Drug Abuse– Intramural Research Program (Z1A DA000606 to L.S., Z1A DA000487 to CRL, Z1A DA000523 to MHB). This work utilized the computational resources of the NIH HPC Biowulf cluster (http://hpc.nih.gov).

## SUPPORTING INFORMATION

**Supplementary Table 1.** Contact frequency and sidechain dihedral angles for CB1R bound with 5F-M<u>D</u>MB-<u>P</u>ICA (DP), 5F-M<u>M</u>B-<u>P</u>ICA (MP), and M<u>D</u>MB-<u>F</u>UBINACA (DF) Residues are shown as one letter with residue numbers and Ballesteros-Weinstein numbering for GPCRs [83] in parenthesis. We used 25% difference as the threshold to identify residues having divergent contact frequencies and 60-degree difference as the threshold to identify the residues having divergent sidechain rotamers (or the distributions).

| Residue | Contact Frequency |  |  |  |  | PIA-GPCR (Sidechain dihedral angles) |  |  |  |  |
| --- | --- | --- | --- | --- | --- | --- | --- | --- | --- | --- |
|  | DP | MP | DF | (DP – MP) | (DF – MP) | DP | MP | DF | (DP – MP) | (DF – MP) |
| I169 (2.56) | 43.6 | 93.4 | 66.0 | -49.7 | -27.3 | 275.3 | 279.8 | 286.9 | 4.5 | 7.1 |
| F170 (2.57) | 99.9 | 100.0 | 100.0 | 0.0 | 0.0 | 180.7 | 180.3 | 178.9 | 0.4 | 1.4 |
| V171 (2.58) | 0.0 | 66.0 | 20.5 | -66.0 | -45.5 | 233.0 | 203.4 | 212.9 | 29.6 | 9.5 |
| S173 (2.60) | 99.9 | 100.0 | 100.0 | 0.0 | 0.0 | 133.2 | 205.5 | 111.8 | -72.3 | -93.7 |
| F174 (2.61) | 99.9 | 98.6 | 92.7 | 1.3 | 6.0 | 200.1 | 185.3 | 190.7 | 14.8 | 5.4 |
| F177 (2.64) | 99.9 | 99.9 | 100.0 | 0.0 | 0.0 | 187.2 | 247.8 | 228.2 | 60.6 | 19.6 |
| H178 (2.65) | 95.4 | 46.3 | 100.0 | 49.1 | 53.6 | 276.1 | 285.0 | 258.6 | 8.9 | 26.4 |
| F189 (3.25) | 89.6 | 96.9 | 99.8 | 7.3 | 2.8 | 178.5 | 180.7 | 179.6 | 2.2 | 1.1 |
| K192 (3.28) | 67.6 | 94.2 | 90.1 | 26.6 | 4.1 | 281.9 | 286.1 | 288.1 | 4.2 | 2.0 |
| L193 (3.29) | 99.9 | 100.0 | 100.0 | 0.0 | 0.0 | 191.9 | 181.0 | 185.1 | 10.9 | 4.1 |
| G194 (3.30) | 0.0 | 60.4 | 0.0 | -60.4 | -60.4 | 0.0 | 0.0 | 0.0 | 0.0 | 0.0 |
| V196 (3.32) | 99.9 | 100.0 | 100.0 | 0.0 | 0.0 | 178.8 | 169.3 | 180.5 | 9.5 | 11.2 |
| T197 (3.33) | 99.9 | 100.0 | 100.0 | 0.0 | 0.0 | 308.3 | 306.6 | 307.4 | 1.7 | 0.8 |
| F200 (3.36) | 99.9 | 99.6 | 99.5 | 0.3 | 0.1 | 278.4 | 274.3 | 282.2 | 4.1 | 7.9 |
| I267 (ECL2) | 76.0 | 28.5 | 55.3 | 47.5 | 26.8 | 267.8 | 206.9 | 262.3 | 60.9 | 55.4 |
| F268 (ECL2) | 99.9 | 100.0 | 100.0 | 0.0 | 0.0 | 289.9 | 279.8 | 288.3 | 10.1 | 8.5 |
| P269 (ECL2) | 60.4 | 0.0 | 10.1 | 60.4 | 10.1 | 166.5 | 89.4 | 207.8 | 77.1 | 118.4 |
| H270 (ECL2) | 43.3 | 16.8 | 50.0 | 26.5 | 33.2 | 221.1 | 240.2 | 150.1 | 19.1 | 90.1 |
| I271 (ECL2) | 98.3 | 82.8 | 99.0 | 15.5 | 16.2 | 131.8 | 145.0 | 130.0 | 13.2 | 15.0 |
| D272 (5.36) | 29.6 | 11.5 | 53.1 | 18.2 | 41.6 | 183.1 | 183.5 | 183.1 | 0.4 | 0.4 |
| Y275 (5.39) | 99.8 | 88.4 | 100.0 | 11.3 | 11.5 | 178.5 | 179.9 | 180.8 | 1.4 | 0.9 |
| L276 (5.40) | 99.8 | 96.0 | 100.0 | 3.9 | 4.0 | 217.3 | 254.5 | 218.8 | 37.2 | 35.7 |
| W279 (5.43) | 99.8 | 100.0 | 100.0 | 0.2 | 0.0 | 185.9 | 185.8 | 186.4 | 0.1 | 0.6 |
| W356 (6.48) | 62.2 | 57.3 | 69.3 | 5.0 | 12.0 | 293.5 | 293.5 | 295.4 | 0.0 | 1.9 |
| L359 (6.51) | 99.4 | 99.4 | 100.0 | 0.0 | 0.5 | 227.2 | 250.2 | 238.3 | 23.0 | 11.9 |
| M363 (6.55) | 99.9 | 99.4 | 99.9 | 0.5 | 0.5 | 287.1 | 278.0 | 289.4 | 9.1 | 11.4 |
| F379 (7.35) | 99.5 | 97.6 | 98.4 | 1.9 | 0.8 | 178.4 | 181.5 | 181.9 | 3.1 | 0.4 |
| C382 (7.38) | 31.0 | 51.4 | 30.6 | 20.4 | 20.8 | 287.3 | 290.2 | 287.6 | 2.9 | 2.6 |
| S383 (7.39) | 99.9 | 98.5 | 100.0 | 1.4 | 1.5 | 182.8 | 137.4 | 213.8 | 45.4 | 76.4 |
| C386 (7.42) | 98.9 | 94.7 | 99.6 | 4.1 | 4.9 | 286.2 | 286.2 | 295.0 | 0.0 | 8.8 |

**Supplementary Figure 1.**
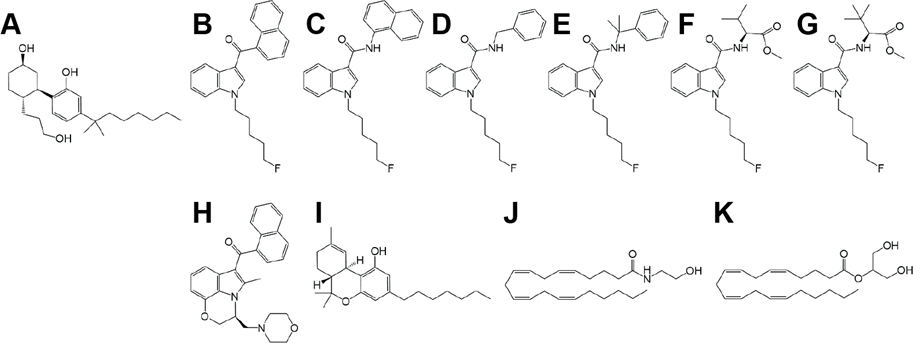
Chemical structures. A. CP55940, B. AM2201, C. 5F-NNEI, D. 5F-SDB-006, E. 5F-CUMYL-PICA, F. 5F-MMB- PICA, G. 5F-MDMB-PICA, H. WIN55-212-2, I. Δ^9^-THC, J. AEA, K. 2-AG

**Supplementary Figure 2.**
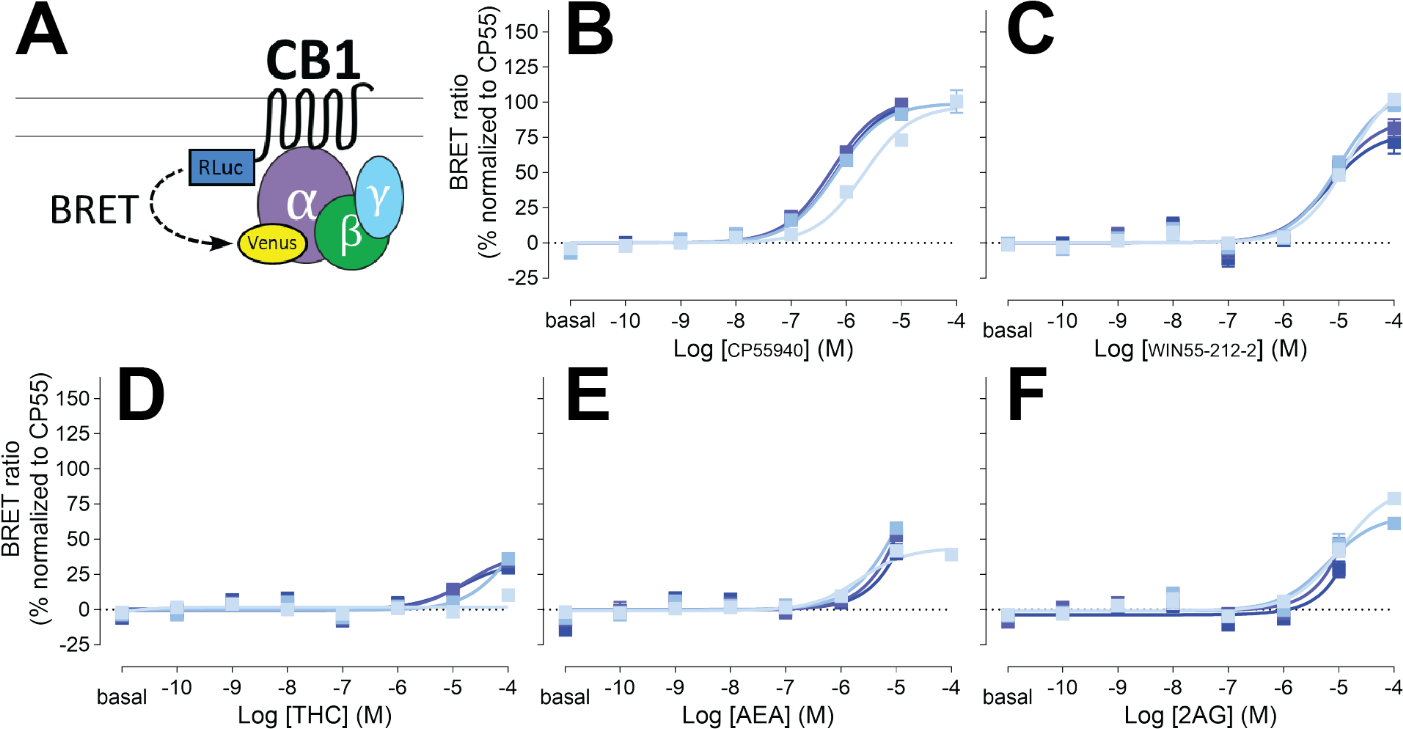
G_i1_ engagement BRET. Drug-induced engagement BRET between CB1R-Rluc and Gi1-Venus (A) measured in response to CP55940 (B), WIN55-212-2 (C), Δ^9^-THC (D), anandamide (E), 2- arachidonoylglycerol (F) at various time points (2, 16, 30, 44 min light to dark blue). Concentration-response curves are plotted as a percentage of maximal response by CP55940 at each time point and presented as means ± SEM of *n* ≥ 3 independent experiments.

**Supplementary Figure 3.**
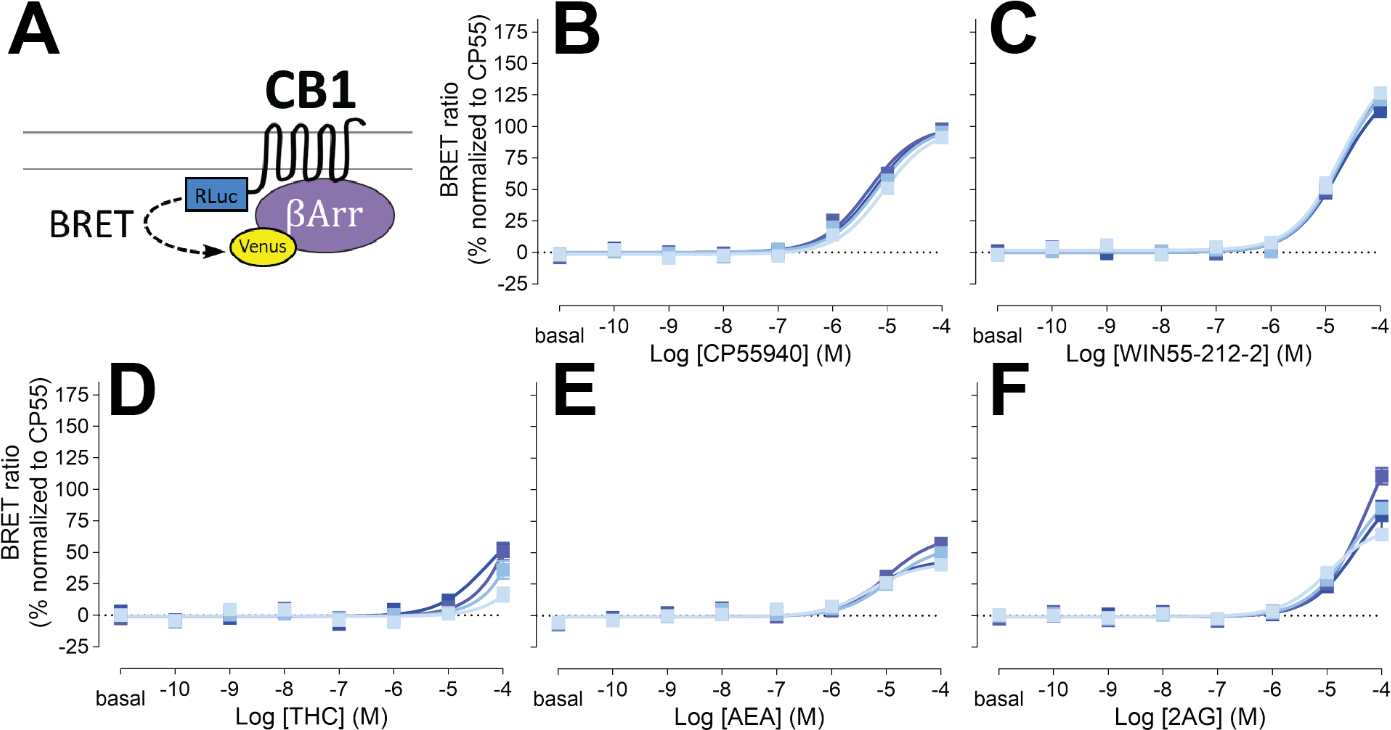
β-arrestin 2 recruitment BRET. Drug-induced recruitment BRET between CB1R-Rluc and β-arrestin 2-Venus (A) measured in response to CP55940 (B), WIN55-212-2 (C), Δ^9^-THC (D), anandamide (E), 2- arachidonoylglycerol (F) at various time points (2, 16, 30, 44 min light to dark blue). Concentration-response curves are plotted as a percentage of maximal response by CP55940 at each time point and presented as means ± SEM of *n* ≥ 3 independent experiments.

**Supplementary Figure 4.**
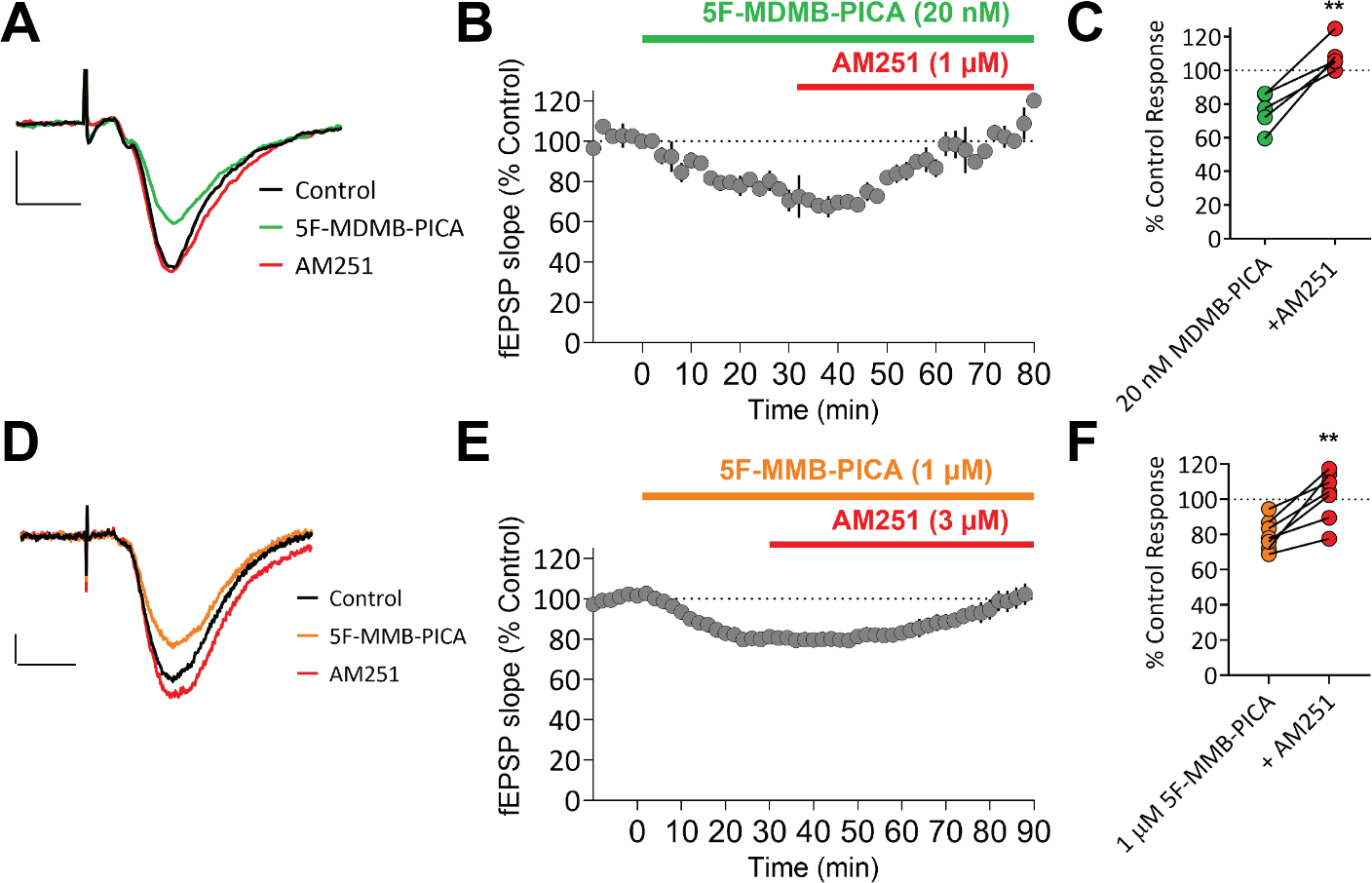
Brain slice electrophysiology. Reversal of hippocampal glutamate release inhibition by CB1R selective antagonist. (A,D) Representative averaged traces for 5F-MDMB-PICA and its reversal by AM251 (A) and 5F- MMB-PICA and its reversal by AM251 (D), (B,E) Summary time course of recordings (*n* ≥ 3 recordings) demonstrating the effects of 5F-MDMB-PICA (B), 5F-MMB-PICA (E), and their reversal. (C,F) Individual recording result of reversal effect by AM251 for 5F-MDMB-PICA (C) and 5F-MMB-PICA (F). Data points are presented as means ± SEM. Paired t-test shows **p < 0.01.

**Supplementary Figure 5.**
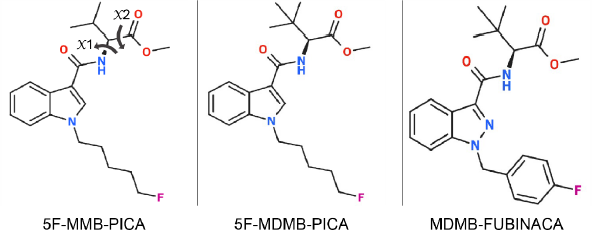
2D structure of 5F-MMB-PICA, 5F-MDMB-PICA, and MDMB-FUBINACA. The head moiety dihedral angles (*x*1 and *x*2) are shown as arrows which are rotation around N-C and C-C, respectively. The same dihedral angle definitions apply to 5F-MDMB-PICA and MDMB-FUBINACA.

**Supplementary Figure 6.**
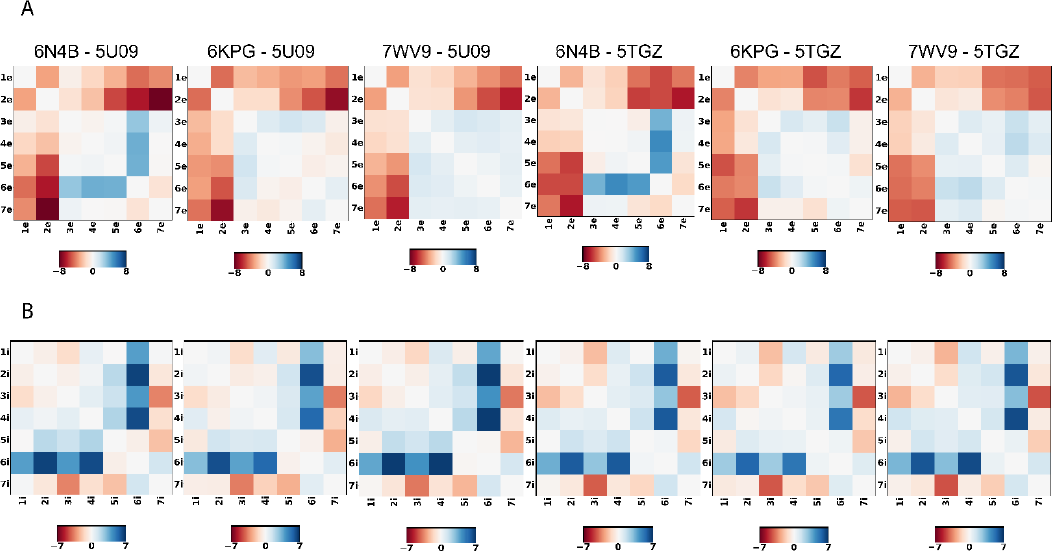
PIA-GPCR analysis. The comparisons are between active hCB1-Gi state (PDBs: 6N4B, 6KPG, and 7WV9) and inactive hCB1 (PDB: 5U09 and 5TGZ). (A) The COM distances between extracellular domain of TMs denoted as (1-7)e. (B) The COM distances between the intracellular domain of TMs denoted as (1-7)i. Among the three active structures, 6N4B has the larger differences from the inactive structures on the extracellular side, which is represented by the larger inward movement of TM2e and outward movement of TM6e. On the intracellular side, due to the conforming effect of binding to the Gi protein, no drastic difference was observed.

